# Prolactin maintains parental responses and alters reproductive axis gene expression, but not courtship behaviors, in both sexes of a biparental bird

**DOI:** 10.1101/2021.12.13.472470

**Authors:** Victoria S. Farrar, Laura Flores, Rechelle C. Viernes, Laura Ornelas Pereira, Susan Mushtari, Rebecca M. Calisi

## Abstract

Prolactin, a hormone involved in vertebrate parental care, is hypothesized to inhibit reproductive hypothalamic-pituitary-gonadal (HPG) axis activity during parenting, thus maintaining investment in the current brood as opposed to new reproductive efforts. While prolactin underlies many parental behaviors in birds, its effects on other reproductive behaviors, such as courtship, remain unstudied. How prolactin affects neuropeptide and hormone receptor expression across the avian HPG axis also remains unknown. To address these questions, we administered ovine prolactin (oPRL) or a vehicle control to both sexes in experienced pairs of the biparental rock dove (*Columba livia*), after nest removal at the end of incubation. We found that oPRL promoted parental responses to novel chicks and stimulated crop growth compared to controls, consistent with other studies. However, we found that neither courtship behaviors, copulation rates nor pair maintenance differed with oPRL treatment. Across the HPG, we found oPRL had little effect on gene expression in hypothalamic nuclei, but increased expression of *FSHB* and hypothalamic hormone receptor genes in the pituitary. In the gonads, oPRL increased testes size and gonadotropin receptor expression, but did not affect ovarian state or small white follicle gene expression. However, the oviducts of oPRL-treated females were smaller and had lower estrogen receptor expression compared with controls. Our results highlight that some species, especially those that show multiple brooding, may be able to maintain mating behavior despite elevated prolactin. Thus, mechanisms may exist for prolactin to promote investment in parental care without concurrent inhibition of reproductive function or HPG axis activity.

## Introduction

Animals that exhibit parental care must balance current reproductive efforts that prioritize care and provisioning of the current brood versus future reproductive opportunities, such as mating (Stearns, 1976). These reproductive transitions require physiological mediators (Ricklefs and Wikelski, 2002; Zera and Harshman, 2001), including endocrine mechanisms that are well-known to facilitate investment and resources in line with such life-history tradeoffs (Hau and Wingfield, 2011; Williams, 2012).

Prolactin, a hormone involved in parental care across vertebrates (Bachelot and Binart, 2007; Freeman et al., 2000), can mediate key transitions in reproductive investment and behavior. Prolactin (PRL) is a 23 kD peptide protein produced in the anterior pituitary with receptors expressed in nearly every tissue across the body, including the brain (Grattan and Bridges, 2009) and gonads (Harris et al., 2004). Best known for facilitating mammalian lactation, PRL also promotes parental motivation and care behaviors in both males and females of many vertebrates (Brown et al., 2017; Hashemian et al., 2016; Smiley, 2019). In order to facilitate reproductive transitions, PRL must interact with the hypothalamus-pituitary-gonadal (HPG) endocrine axis, which coordinates reproductive behaviors and physiology. In the HPG axis, kisspeptin in the hypothalamus stimulates gonadotropin-releasing hormone (GnRH) release onto the anteriory pituitary gland, which then releases luteinizing hormone (LH) and follicle-stimulating hormone (FSH), stimulating the gonads to release sex steroids (e.g. progesterone, testosterone, estradiol).

Additionally, gonadotropin-inhibitory hormone (GnIH) from the hypothalamus can inhibit the release of GnRH, LH and FSH in some vertebrates (Ubuka et al., 2013).

PRL is hypothesized to exert an “anti-gondal” effect on the HPG axis in multiple species. Lactational amenorrhea presents a classic example of this effect, where PRL inhibits ovulation during the energy-intensive period of milk production after pregnancy in mammals (Fourman and Fazeli, 2015). This anovulatory effect is mediated through the inhibition of kisspeptin in rodents (Brown et al 2019, Grattan 2018), which consequently reduces GnRH, gonadotropin and sex steroid release. In seasonally- breeding birds, PRL has been implicated in gonadal regression as birds transition into photorefractoriness (Dawson and Sharp, 1998; Sharp et al., 1998; Small et al., 2008), and a rich body of classic studies show exogenous PRL treatment can induce gonadal regression in birds (Bates et al., 1937; Meier, 1969; Tewary et al., 1983). Further, experiments in doves show that systemic and intracerebroventricular (icv) PRL injections, at levels akin to those circulating during parental care, maintain parental responses and reduce gonad size and LH plasma levels (Buntin et al., 1991, 1988; Buntin and Tesch, 1985; Janik and Buntin, 1985). The evidence for an inhibitory effect of PRL on the HPG axis has been connected to its possible role in photorefractoriness (Dawson and Sharp, 1998), clutch size regulation (Ryan et al., 2015; Sockman et al., 2000) and parental care (Angelier et al., 2016; Buntin, 1996).

In contrast, many other studies have not found evidence for such an anti-gonadal effect of PRL. For instance, classic studies in multiple bird species did not find any effect of PRL treatment on gonadal regression (house finches- Hamner 1968; quail - Jones 1969; sparrow *spp.* - Meier and Dusseau 1968, Laws and Farner 1960). Although an anti-gonadal effect seems clear in male doves, not all studies found changes in testes weight or LH with icv PRL treatment (Foreman et al., 1990). In domestic fowl, immunization against PRL, which reduces functional peptide levels, did not affect LH levels (Crisóstomo et al., 1998), and reduced laying rates (Li et al.,2011). Further, a pro-gonadal function appears in some seasonally-breeding mammals, such as hamsters and sheep, where PRL administration can actually *stimulate* testes growth in periods of gonadal recrudescence (Bartke et al., 1980, 1975; Howell-Skalla et al., 2000; Sanford and Baker, 2010). This mixed literature indicates that PRL’s effects on the HPG axis may be species and breeding context- specific.

Despite this mixed evidence, few studies have examined how non-parental reproductive behaviors respond to PRL, and how multiple HPG signals and receptors may mediate this relationship. Many studies have established PRL’s role in parental behaviors (Brown et al., 2017; Buntin et al., 1991; Horseman and Buntin, 1995), but its influence, if any, on other reproductive behaviors, like courtship and mating, remains understudied. Most of these studies measured gonad size, and/or plasma hormone levels, such as LH or sex steroids. However, hormone receptor expression, which modulates tissue responsiveness to circulating signals, may also underlie any effect PRL has on the HPG axis. Further, no study to our knowledge has examined if there is a causal relationship between gonadotropin inhibitory hormone (GnIH, also known as RFRP-3) and PRL. GnIH has been shown to increase during transitions in parental care in starlings and rodents (Calisi et al., 2016), including at timepoints where PRL levels also rise in birds (Austin et al., 2021; Dawson and Sharp, 1998). Thus, GnIH and its receptor may be an important, but unexplored, HPG target of PRL.

To address these questions, we manipulated PRL levels and examined the effects on HPG axis gene expression and non-parental reproductive behaviors in a monogamous, biparental bird, the rock dove (*Columba livia*). Both male and female rock doves participate in all stages of offspring care, including nest building, egg incubation and chick provisioning (Abs, 1983). Both sexes also produce crop milk, a nutrient-rich substance which offspring depend exclusively on for the first few days of life (Davies, 1939). Like mammalian milk, crop milk production is also driven by PRL (Horseman and Buntin 1995). This avian model thus removes the confounding effects of female-only pregnancy and lactation seen in mammals, allowing us to compare sex-typical biases in HPG regulation and behavior. Lastly, we capitalized upon detailed ethograms of courtship behaviors developed for doves to quantify species-specific reproductive behaviors (Cheng, 1973; Goodwin, 1983, 1956; Lehrman, 1964).

Our study had three main aims: to examine the effect of PRL on 1) parental responsiveness, 2) non-parental reproductive behaviors, like courtship and copulation, and 3) gene expression of hormones and their receptors across the entire HPG axis, with a specific eye towards effects on GnIH. We administered exogenous ovine prolactin (oPRL) or a vehicle control to experienced breeding pairs of rock doves and then removed their nests, forcing birds from a parental state back to a courtship/mating state as they restart their nest efforts. We then recorded parental behaviors in response to a novel chick and observed naturally-occurring courtship behaviors. We also collected brain, pituitary and gonadal tissues to measure HPG gene expression. We hypothesized that oPRL would maintain the HPG in a parental state, favoring current reproductive effort and depressing gonadal activity that may promote future reproductive efforts. Thus, we predicted that oPRL-treated birds would exhibit more parental behaviors in response to a novel chick, would be more delayed in the progression of the dove courtship cycle, and would copulate less often than vehicle-treated pairs. We also predicted that oPRL treatment would reduce GnRH, LH and FSH expression in the hypothalamus and pituitary, increase GnIH expression, and alter gonadotropin receptor expression in the gonads.

## Methods

### Hormone manipulation

To determine how PRL may causally drive transitions between parental and other reproductive behaviors, we treated breeding pairs of rock doves (*Columba livia*) with exogenous PRL during a transition where birds are switching between parental and mating behavior. On day 16 of the 17 day incubation period, we surgically implanted osmotic pumps (Model 2001, release rate: 1.0 µl/hr, Alzet, DURECT Corp.) containing either ovine PRL in 0.87% physiological saline (dose of 3.33 µg/hr, 80 µg/day; A.F. Parlow, National Hormone and Peptide Program) or vehicle (0.9% physiological saline;1.0 µl/hr, 24 µl/day) in both the male and female of the breeding pair. We did not measure circulating PRL because oPRL is not reliably detected by chicken PRL antibodies, and endogenous PRL would likely be lower in oPRL-treated birds due to negative feedback (Z.Wang, F. Angelier *pers. comm.*). However, using the plasma concentration estimation described in Sockman et al., (2000), we estimate that this dose will lead to a plasma concentration of approximately 46.25 ng/mL in the average rock dove in our population. This concentration is on par with PRL levels observed in late incubation and during nestling rearing in previous studies of rock doves (Austin et al., 2021). These osmotic pumps reliably release 1.0 µl/hr for seven days (Alzet, DURECT Corp.), thus the experimental period lasted seven days (Figure 1). oPRL delivered through osmotic pumps has been shown to activate PRL receptors and signalling pathways in the brain of a closely-related species, the ring dove (Buntin and Buntin, 2014), and we replicated the osmotic pump model and dose used in that study.

**Figure 1.**
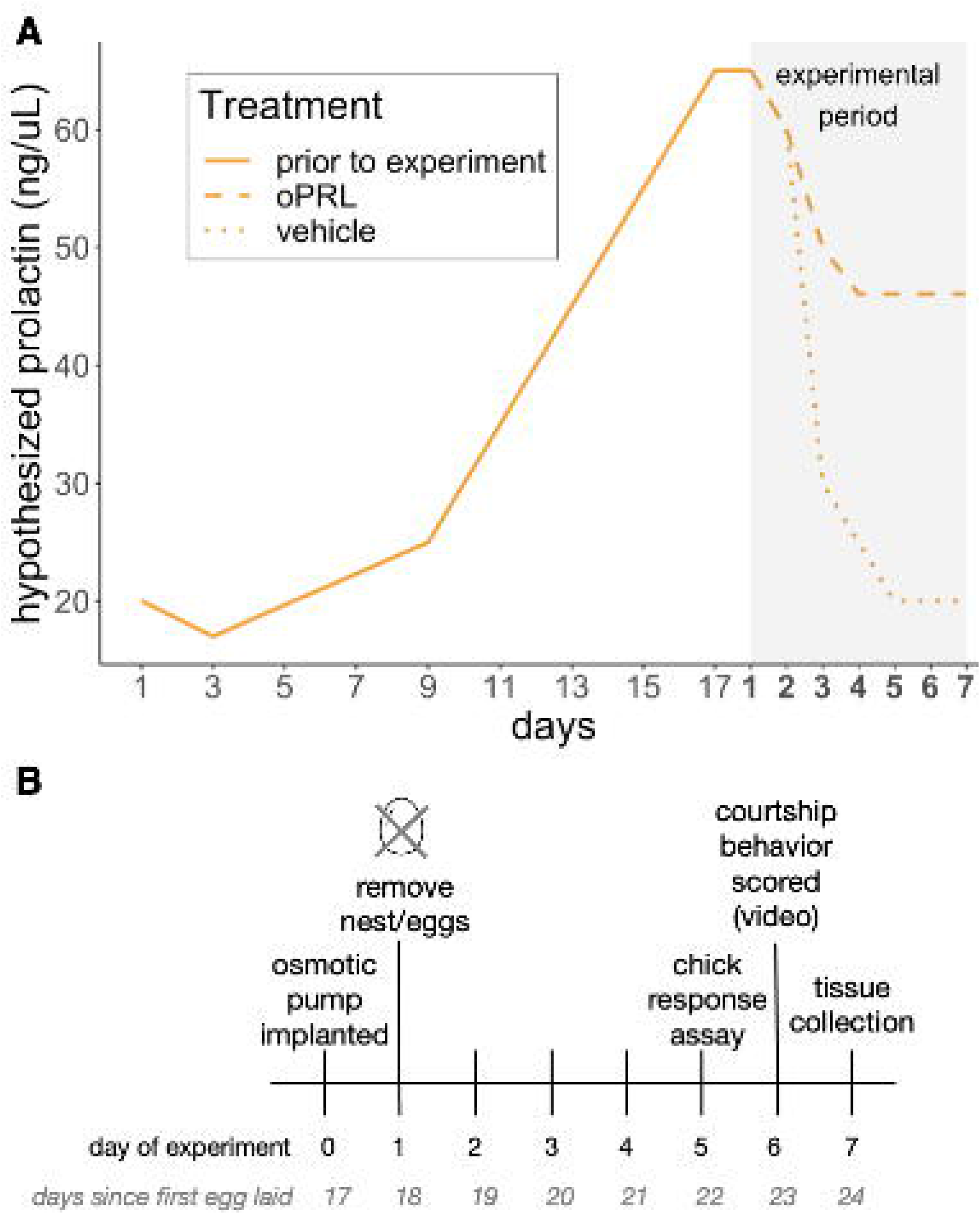
Experimental timeline. **(A)** Hypothesized concentrations of prolactin during the typical dove incubation period and as expected during the experimental period, both in the experimental treatment (oPRL, dashed) and control (vehicle, dotted) groups. Hormonal data during incubation based on data from Austin et al., 2021; Dong et al., 2012 and expected experimental values are drawn from patterns seen in ring doves (Lea & Sharp, 1991; Ramsey et al., 1985). The experimental period is shaded in gray. **(B)** Both male and female birds in a pair had osmotic pumps containing either ovine prolactin or vehicle implanted surgically on day 17 of incubation. The experiment began on the expected hatch date, incubation day 18, when eggs and nests were removed, and ended seven days later when tissues were collected. For full details, see Methods.

Sixteen breeding pairs received pumps with oPRL (32 birds), while 12 received vehicle (24 birds), for a total of 56 birds. All breeding pairs used in this study were reproductively experienced and had raised at least one chick to fledging before the experiment. Birds were socially housed in a semi-natural environment, where they selected their own mate and nest box within a colonial aviary, which houses 10 - 12 other breeding pairs (see MacManes et al., 2017 for details).

To induce a shift from parental behaviors to re-mating, we removed eggs and hay from the nest box on incubation day 17, the day after surgical pump implantation and the last day of incubation before expected hatching (hatching typically occurs after 18 days of incubation). Eggs and hay were removed between 09:00-11:00 hours on incubation day 17. As rock doves breed nearly continuously in captivity (Johnston, 1992), egg removal ends the current nest effort and birds will court and re-mate to start a new clutch.

### Behavior: Response to novel chick

As a proof of principle that PRL promotes parental behaviors and responses, we tested birds in both groups on their response to a novel chick after nest loss. On the fifth day of the seven-day experimental period (Fig.1), we placed a young chick (average age: 6.6 days old) from another nest into the focal pair’s nest box for approximately 120 minutes between 08:00 - 12:00, and video-recorded the behavioral response. To ensure that chicks elicited parental responses from subjects, we removed chicks from their home nests at least twelve hours before the behavioral trial to restrict them from feedings and induce begging behavior when placed in the subject’s nest. Unlike previous studies in ring doves (Buntin et al., 1991; Wang and Buntin, 1999), we did not remove either sex during the behavioral trial, as to avoid introducing a stressor (due to loss of the mate or separation from the mate, or the stress of being isolated while the mate was tested). Instead, we recorded the response of both birds in the pair simultaneously during these trials. At the end of the live behavioral trial, we measured the change in weight of the chick compared to at the start of the trial (post-trial weight minus pre-trial weight / pre-trial weight), as an additional measure of feeding attempts by the focal subjects.

Video recordings were scored on BORIS v.7.9 (Friard and Gamba, 2016) by trained observers who were blind to treatment. We recorded the occurrence and duration of the following chick-oriented behaviors: a) entering and standing in the nest box while the chick is present, b) brooding the chick, by standing or squatting over the chick, with body-to-body contact, and c) feeding the chick, where a bird engaged in mouth-to-mouth regurgitation behaviors with the chick. We also recorded the duration of time spent aggressing the chick (vigorously, offensive pecking) if it occurred. For each pair, we averaged the time each bird spent expressing each behavior during the trial, and present this data as a percent of the total trial time recorded. Two trials were excluded from behavioral scoring because the video recording equipment failed to record or save the recording. Both of these excluded trials came from oPRL-treated pairs, but we still had chick weight data as a proxy for feeding attempts by these pairs. In total, we scored videos for 26 pairs (n = 14 oPRL, n = 12 vehicle).

To analyse these behavioral responses, we first compared the percent of pairs that expressed each of the four behaviors across treatment groups (i.e. oPRL vs vehicle) using pairwise chi-square tests. Then, we compared whether the percent of time pairs expressed each behavior differed with treatment using pairwise t-tests.

### Behavior: Courtship and reproductive behaviors

To examine how PRL may affect non-parental reproductive behaviors, we recorded occurrences of various courtship behaviors and copulations for a pre-selected subset of pairs. On day six of the seven day experiment (Fig.1), we recorded video from two cameras, one placed facing the focal pair’s nest box and one recording activity on the entire aviary cage floor. We recorded reproductive behaviors on day six of the seven day experimental period for two reasons. First, this timepoint was furthest in time from nest removal (and thus closest to any possible ovulation/egg laying event, if it occurs), and second, these behaviors would be the most temporally connected to gene expression as tissues were collected the morning of day seven. We recorded videos for eight oPRL-treated pairs (n = 16 birds) and seven vehicle-treated pairs (n = 14 birds) in total.

Nest box videos were used to measure species-specific courtship behaviors observed in doves, such as bow-coos, nest-coos, and nest building through hay manipulation (Cheng, 1973). Nest videos were recorded from 07:30 to 18:30 for all pairs, and 30-minute videos for scoring were sampled every 90 minutes, resulting in four hours (240 minutes) of total scored nest video per pair. Trained observers, blinded to treatment, scored videos in BORIS v.7.9 (Friard and Gamba, 2016) for the following dove courtship behaviors: a) bow-cooing in the nest, where the bird struts and turns in the nest box while cooing and stamping feet up and down (Goodwin, 1956)**, b)** nest-cooing or “nodding”, where the bird takes a posture with head down and tail up and inflates the crop to coo (Cheng, 1973; Goodwin, 1956), c) wing-flipping, where the bird’s head is low, tail is up and it gently flips or flicks the tips of its wings while nest-cooing (note: this behavior always occurs during nest cooing, but nest cooing can occur without wing-flipping Cheng, 1973; Miller and Miller, 1958), and d) nest-building, via hay manipulation, where a bird picks up a single hay stick into its bill, with or without bringing the hay to the nest (Cheng and Balthazart, 1982; Miller and Miller, 1958). These courtship behaviors have been well-described in the stereotypical courtship progression of doves, including *Columba livia* (Cheng, 1992; Goodwin, 1983).

We also scored the following pair-maintenance behaviors: a) allopreening or “hetero-preening”, where the bird preens its mate, typically around the head and neck (Miller and Miller, 1958), b) allofeeding or billing, where a bird, typically the male, opens their bill towards the mate and feeds the mate similarly to how chicks would be fed (this usually occurs after a bout of allopreening) (Miller and Miller, 1958). These pair maintenance behaviors usually precede copulations, for which we scored the following: c) soliciting copulations, where the bird (typically female) squats down, slightly spreads wings and shoulders to facilitate a mount (also called the “sex crouch” - Miller and Miller 1958), and d) mounting, where a bird mounts another in a copulatory position (typically the male) (Goodwin, 1956; Miller and Miller, 1958). All behavioral descriptions are consistent with Miller and Miller 1958, and were aligned to the representative courtship stage as described in Cheng 1992. All behaviors were scored as state events, with a start, end, and duration, except: nest building (hay manipulation), soliciting copulation and mounting, which were scored as point behaviors with only occurrences counted.

As most copulations occur outside the nest box (*pers.obs.*), we also scored aviary floor videos for copulation attempts. Aviary floor videos, which had a lower resolution, were recorded from 07:30 to 18:30 for all pairs, and all 11 hours of the floor video was scored to capture the daily copulation rate. Copulation attempts were scored by counting the occurrence of a) soliciting copulations, and b) mounting, as described above. Interestingly, reciprocal mountings, where males appear to “solicit” mountings by females, or females mount their mate, have been observed in rock doves (Goodwin, 1956), and we observed both males and females mounting and soliciting. However, we only compared male mounts and female soliciting in our data analyses.

To analyse courtship and other reproductive behaviors, we sorted behaviors by the sex that performs them typically during the stereotyped dove courtship cycle (Cheng, 1992). We compared the following behavior*sex combinations: male bow-coo, male nest-coo and wing-flip, female nest-coo and wing-flip, male billing, male mounting,and female soliciting copulations. We compared allopreening and nest-building in both sexes. For each behavior*sex combination, we then compared if the proportion of time spent exhibiting the behavior or occurrence (for state and point behaviors, respectively) differed between oPRL and vehicle-treated birds using pairwise t-tests.

### Tissue Collection

Seven days after surgery (six days after nest removal), we euthanized birds using an overdose of isoflurane anesthetic followed by swift decapitation. Brains were removed and flash frozen on dry ice within 3 minutes of capture from the cage (as described in MacManes et al 2017). Trunk blood, pituitary gland, gonadal tissue, and crop sac tissue were flash-frozen and stored at -80[ for future gene expression analyses. All methods were approved under UC Davis IACUC protocol #20618.

During collection, the entire crop sac was removed from each bird, fat and crop milk removed, patted dry and weighed for wet weight before flash freezing. We also measured the weight of the ovaries and oviduct combined in females and both testes in males, the diameter and state of the largest ovarian follicle in females, and the length of the testes in males. Crop and gonad weights were normalized by dividing by overall bird body weight. We assessed normality of distributions using Shapiro-Wilks tests, and since all but testes length were significantly non-normal (*p* < 0.05), we compared crop and gonad sizes across treatment groups using the non-parametric Mann-Whitney U tests instead of t-tests.

### Hypothalamic nuclei microdissection

To examine expression in specific nuclei of the hypothalamus, we microdissected the paraventricular nucleus (PVN) and preoptic area (POA) with 2 mm punches using a Leica CM1950 cryostat. We used nuclei landmarks described in Karstens & Hodos 1966 to isolate nuclei. Briefly, we started collecting the POA when the tractus septomesencephalicus (TSM) terminates at the bottom of the brain, stopping when the TSM is no longer visible, and began collecting the PVN when the quintofrontal tracts appear until the optic tecta are visible (see Supp. Table 1 for details and Figure 4 for representative punches). Hypothalamic nuclei punches were weighed and then stored in 200 µL of TriSure reagent (BioLine) at -80 until RNA extraction.

### Quantitative PCR

We extracted RNA from the brain nuclei punches, whole pituitary glands, and gonad samples using TriSure (BioLine) along with a modified protocol of the Zymo Direct-zol RNA miniprep spin-column extraction kit (Zymo Scientific). For gonad analysis, we took a 10 mg sample from the midsection of the left testis, a 10 mg sample of small white follicles (pre-yolk deposition) from the ovary, and a 10 mg sample of the magnum region of the oviduct (Apperson et al., 2017). We measured total RNA quality using a Nanodrop ND-1000 spectrophotometer (ThermoScientific) and RNAs with 260/280 and 260/230 ratios > 1.8 were used in downstream analyses.

We then treated total RNA with a second round of DNase treatment using the Quanta Perfecta DNase kit (QuantaBio), and then converted 8 uL of DNase-treated RNA to complementary DNA (cDNA) using qScript cDNA Supermix (QuantaBio). We diluted cDNA 1:5 for quantitative PCR analyses.

We ran qPCR reactions for the following genes of interest: *GNRH1, GNIHR* in the POA, *GNIH* in the PVN, *AR, ESR1, ESR2, PRLR* in the POA and PVN, *CGA, FSH, GNIHR, GNRHR* in the pituitary, *AROM, FSHR, LHCGR and GNIH* in the testes and ovarian follicles, and *ESR1* and *ESR2* in the oviduct (see Supplemental Table 2 for full gene names and primers). For each tissue, we measured the expression of HPG reproductive hormones or enzymes expressed in that tissue, as well as the relevant receptors to upstream hormones. We measured *GNIH* in the PVN as this is the main location of GnIH-expressing cell bodies in birds (Ubuka et al., 2013). GnIH neurons have been shown to project to GnRH-1 neurons in the POA and the pituitary gland via the median eminence (Ogawa and Parhar, 2014), thus we measured *GNIHR* in the POA and the pituitary gland. As GnIH gene expression has also been measured in avian gonads (Bentley et al., 2017), including the rock dove (MacManes et al., 2017), we also measured GNIH in the testes and follicles. We also included *ACTB, GAPDH, HPRT1, RPL4* as reference genes, which have been shown to be reliable reference genes in avian tissues (Zinzow-Kramer et al., 2014).

We designed all primers used in this study to be specific to *C.livia* using the assembled Rock Dove transcriptome v2.10 (NCBI accession no. GCA_000337935.2). We validated primers by running a 10-fold serial dilution to calculate replication efficiencies (95.6 ± 1.12 %) and confirmed a single product through melt curve analysis (see Supp. Table 2 for accession numbers and primer efficiencies). We ran qPCR reactions in triplicate for each sample using the following mix: 1 µL cDNA template (diluted 1:5), 5 µL 2X SSOAdvanced SYBR Green PCR mix (BioRad), and 10 µM each of primer ( total volume: 10 µL). 384-well qPCR plates were run on a CFX384 Touch Real-Time PCR detection system (BioRad) in the following thermocycling conditions: 50 for 2 min, 95 for 10 min, and then 40 cycles of 95 for 15 sec and 60 for 30 sec. All samples for each tissue-gene combination were run on a single plate.

We calculated relative gene expression using the delta-delta-Ct method (2^- C^;^t^ Livak and Schmittgen, 2001). In this method, gene expression is first normalized to the geometric mean of two reference genes run for each sample (dCt). We used *HPRT1* and *GAPDH* as reference genes for hypothalamic nuclei, *HPRT1* and *RPL4* for pituitary glands,*ACTB* and *GAPDH* for ovarian follicles, and *ACTB* and *RPL4* for oviducts and testes. We confirmed each of these reference gene combinations to be stably expressed in each tissue (no effect of treatment, sex, or their interaction on reference gene Ct, Supplemental Table 3). Normalized expression (dCt) was then expressed relative to average value for a “control” sample group, in this case vehicle-treated females (ddCt). Fold change equals 2^(- ddCt)^. We log-transformed fold changes for analysis.

For each tissue-gene combination, we ran independent general linear models (glms) comparing the effect of treatment (oPRL or vehicle), sex (for hypothalamic nuclei and the pituitary), and their interaction on gene expression. For each gonad type (testes or ovaries), which are unique to one sex, glms only included treatment as an independent variable. We present ANOVA run on these glms. We ran each tissue-gene combination as a separate model because a) the ddCt method does not allow for direct comparisons between genes (Livak and Schmittgen, 2001), and b) each of these genes are regulated by different promoters and transcription factors, and are subject to different tissue-specific regulation, thus we considered their expression independently. Due to the number of linear models run, we adjusted p-values for ANOVA using Benjamini-Hochberg false discovery rate (FDR) correction for multiple comparisons (Benjamini and Hochberg, 1995). All statistical analyses were completed in R (v.4.0.3, R Core Team, 2020).

## Results

### Parental behavior and crop weights

As a confirmation of PRL’s effect on parental behavior, we found that oPRL treatment significantly increased the likelihood of exhibiting parental behaviors towards a novel chick after nest removal (Table 1; Fig.2A). Almost all oPRL-treated pairs brooded or fed the novel chick (Table 1; 92.9% of pairs), while only one pair of vehicle-treated birds (8.3% of pairs) exhibited these responses to the chick. Further, oPRL significantly increased the average proportion of time birds spent brooding, and also led to a suggestive trend towards increased time spent feeding (Table 1; as only one pair of vehicle-treated birds showed feeding, the feeding showed by this pair was lower than the average oPRL treated pair, 1.2% of the trial versus an average of 4.4% of the trial for oPRL pairs). Although we did not separate birds from their mates and pairs were tested together, we also examined the likelhiood of each sex to respond in the trial. Most of the response in oPRL-treated pairs was driven by females, as 12 oPRL females (85.7%) fed the chick at least once, compared to one vehicle female (8.3%) (χ = 13.3, *p* < 0.01). In males, five (35.7%) oPRL-treated males fed the chick at least once, compared with no vehicle-treated males (χ^2^= 5.30, *p* = 0.021). Both treatment groups were equally likely to enter the nest box, presumably to investigate the chick, though oPRL led to significantly more time spent in the nest box. Both groups were equally likely to exhibit aggressive behaviors towards the chick, and the average proportion of time spent showing these behaviors did not significantly differ (Table 1).

**Table 1.**
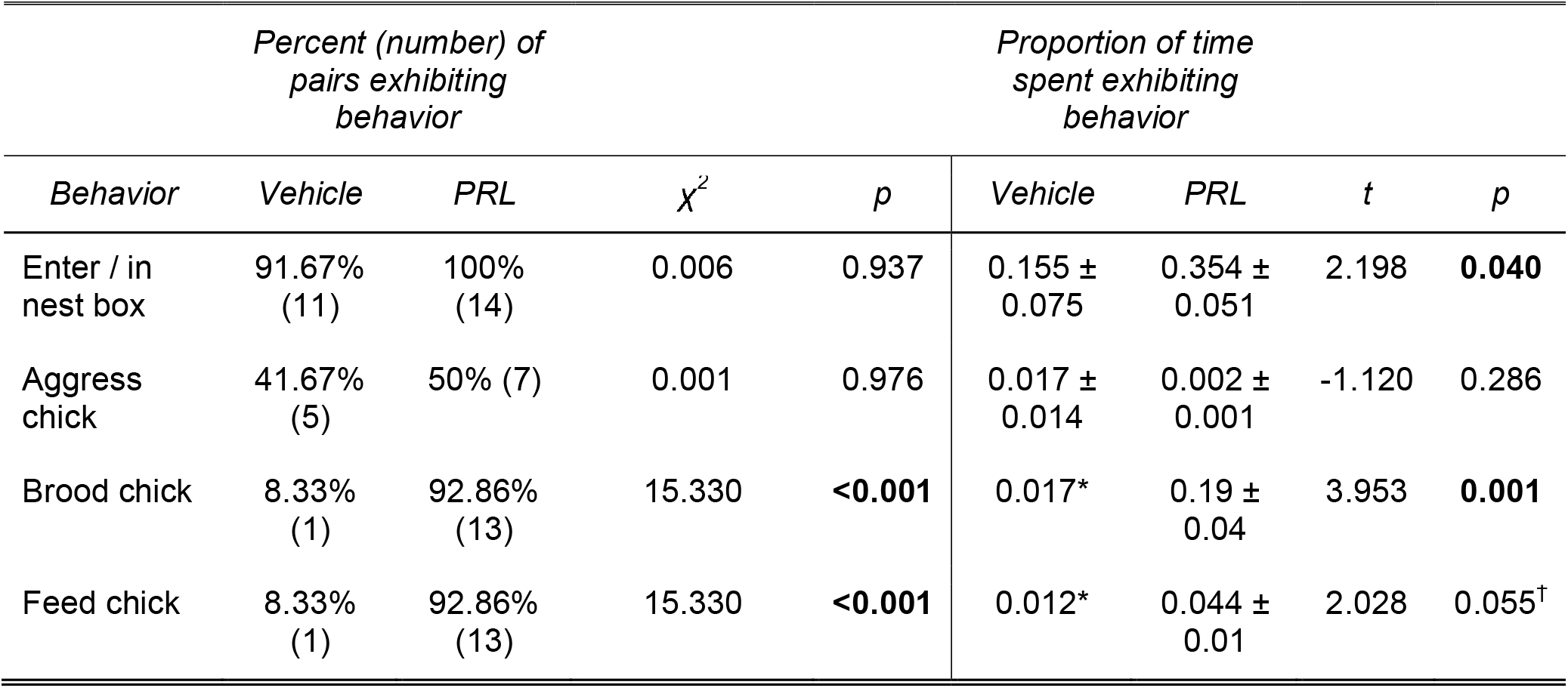
Behavioral responses to novel chick after five days of experimental treatment. The percentage and number of pairs that exhibited the behavior at least once, as well as the average proportion of the assay time (2 hours) those pairs spent exhibiting the behavior are listed. Proportions are listed out of 1, and mean ± SEM are shown. Significant differences between treatment groups (indicated by chi- squared tests for percentage and t-tests for proportions of time) are indicated in bold. *As only one pair exhibited these behaviors, no standard error was calculated. ^†^ represents a suggestive trend towards significance (0.05 < p < 0.10).

**Figure 2.**
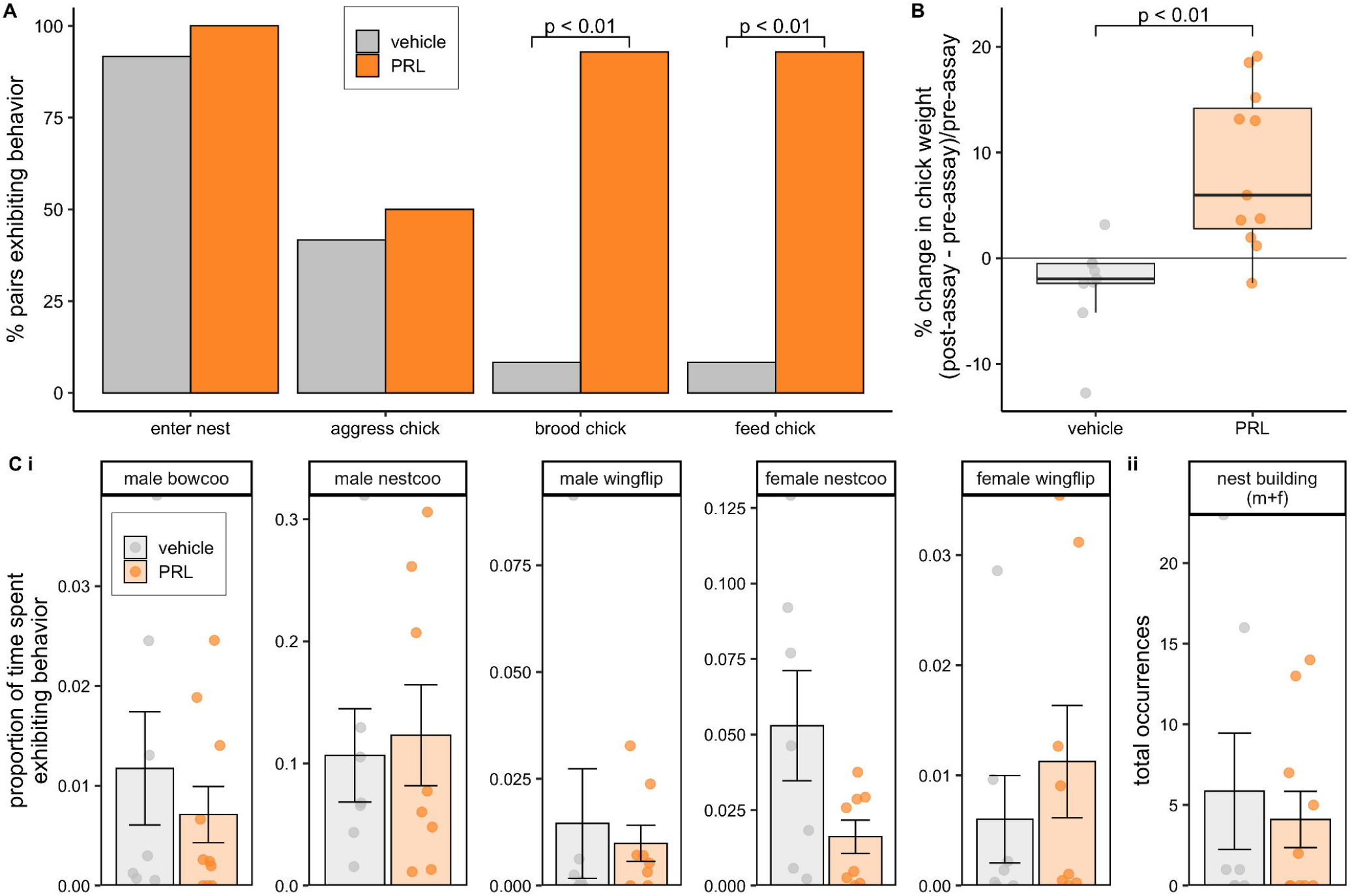
Behavioral responses to a novel chick and observed reproductive behaviors. After five days of experimental treatment (vehicle or oPRL), a novel chick was placed in each pair’s home nest box. (**A**) The percent of pairs tested where at least one bird showed a behavioral response during this chick response assay is shown, as well as (**B**) the percent change in body weight of the stimulus chick, which is measured as post-assay body weight minus pre-assay body weight, divided by the pre-assay body weight. (**C**) After 6 days of treatment, courtship behaviors were observed, with the proportion of the observed time a bird exhibited the behavior (out of 1.0 max, or 100% of the observed time) shown for (**i**) state behaviors and (**ii**) the total number of occurrences shown for point behaviors. Means ± SEM are shown, and points represent pairs (A) or individual birds (B-C).

As further evidence that PRL promoted chick feeding, we found that chicks placed in oPRL nests significantly gained more weight during the trial (Fig.2B; average 9.79% weight increase in oPRL nests vs. 0.08% increase in vehicle nests, Cohen’s *d* = 1.24, *t* = 3.11, *p* = 0.005). When crops were weighed at collection, oPRL-treated birds had significantly larger crops than vehicle-treated birds (Fig.3A; 2.00 % of body weight in oPRL vs 0.68% in vehicle, Cohen’s *d* = 2.81, *t* = 12.20, *p* < 0.001). All oPRL crops had clearly visible crop milk and evidence of thick cornification of the crop (Gillespie et al., 2013) compared to the thinner, less-vascularized crops of the vehicle-treated birds.

**Figure 3.**
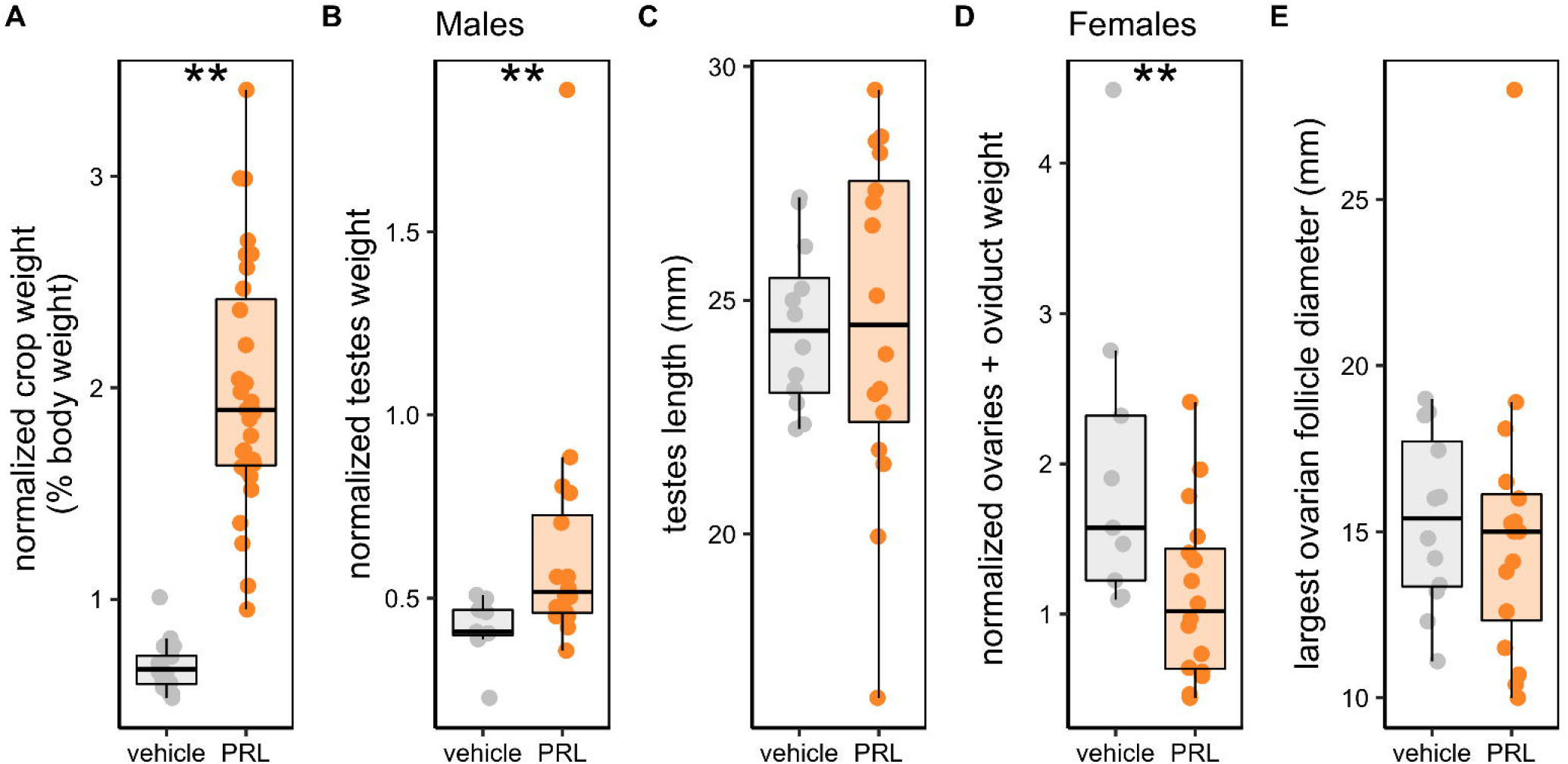
Measurements of crop and gonads in birds treated with oPRL or vehicle. **(A)** Crop sac weights, normalized to body weight (% of overall body weight), (**B**) male testes weight (normalized to body weight), (**C**) vertical length of testes (in millimeters), (**D**), female ovary and oviduct weights, weighed together and normalized to body weight, and **(E)** the diameter (mm) of the largest ovarian follicle (mm) were measured at tissue collection. ** signifies *p* < 0.01 on a Mann-Whitney non-parametric U test. Vehicle-treated birds are shown in gray, oPRL-treated birds in orange. Boxplots show median and first and third quartiles, and dots represent individual samples.

### Courtship and reproductive behaviors

We found no significant effect of treatment on any of the stereotypical dove courtship behaviors (Figure 2C, Table 2; male bow-coo, male nest-coo and wing-flip, female nest-coo and wing-flip, or nest building of either sex, all *p* > 0.09). Further, we found no significant effects of treatment on male or female allopreening bouts, or male-initiated allofeeding/billing, and all birds were observed allopreening their mate at least once (Table 2). We observed male mounting in all pairs. There was no significant effect of treatment on copulation rates, measured either through male mounting or female solicitations (all *p* > 0.1, Table 2).

**Table 2.**
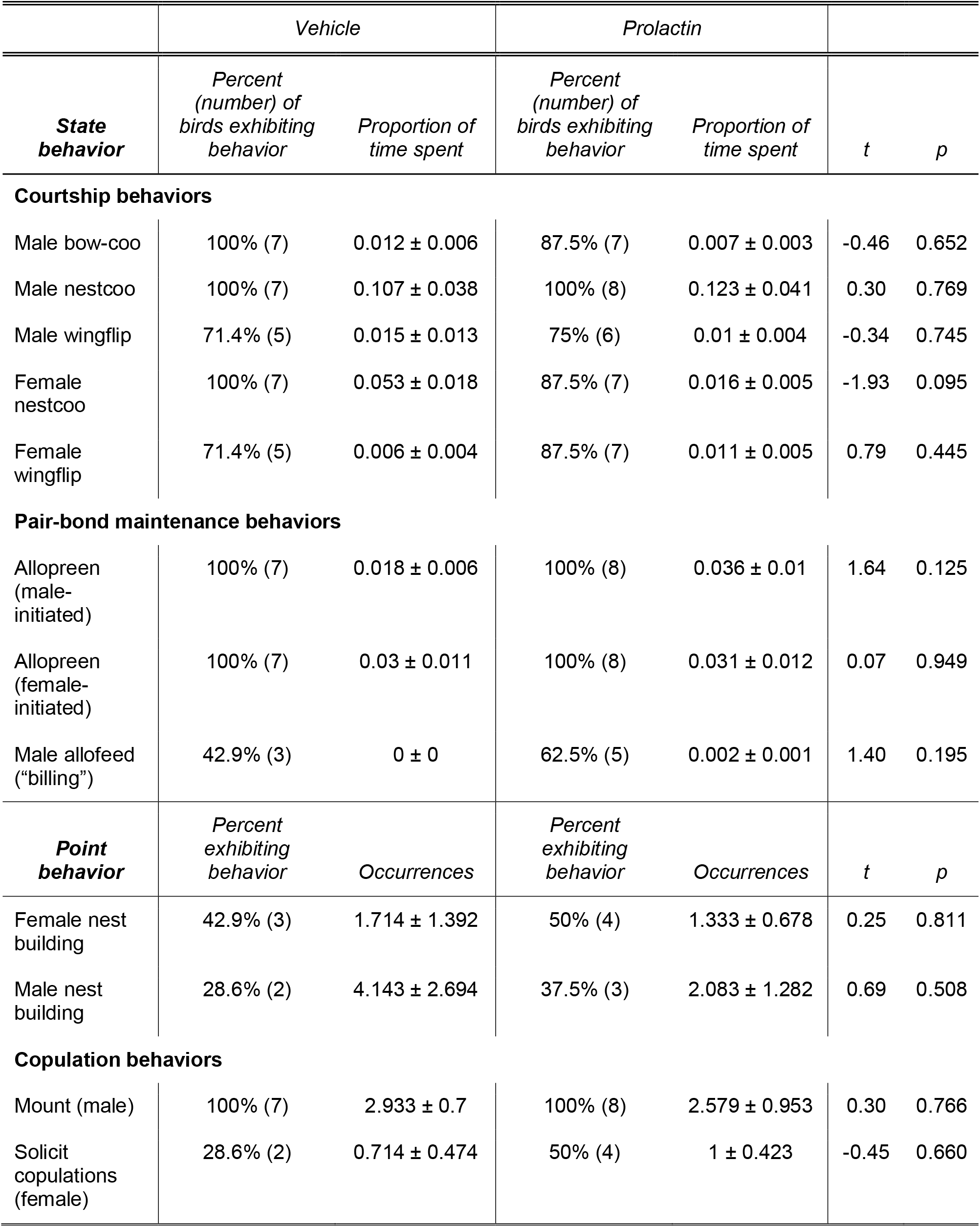
Reproductive Behaviors

### Gonad morphology

We found a significant effect of treatment on both male and female gonad weights. Normalized testes weights were significantly higher in oPRL-treated birds than vehicle birds (*U = 117, p* = 0.010, Cohen’s *d* = 0.76). Female normalized ovaries and oviduct weights were significantly lower in oPRL birds than vehicle birds (*U = 30*, *p* = 0.017, Cohen’s *d* = -1.08). However, the diameter of the largest ovarian follicle did not differ significantly with treatment (*U* = 81.5, *p* = 0.516, Cohen’s *d* = - 0.08); all females had large yolky follicles at collection. Similarly, there was no significant difference in testes length between treatment groups in males (*t* = 0.11, *p* = 0.909, Cohen’s *d* = 0.04). No birds laid eggs during the experimental period, and no females were near laying at collection.

### Hypothalamic gene expression

In the POA, we saw no significant effect of treatment or sex on *GNRH* or *GNIHR* expression (Table 3; Fig.4). Similarly, we saw no significant effect of treatment or sex on *GNIH* expression in the PVN (Table 3; Fig.4C). As for sex steroid receptors measured in both the POA and the PVN, there was no effect of treatment, sex, or nuclei on *AR* expression. *ESR1* tended to be expressed at lower levels in oPRL-treated birds compared to vehicle (*p* = 0.04), but after Benjamini-Hochberg corrections, this effect was no longer significant (*p_adj_* = 0.16; Supplemental Table 4). There was no effect of sex or nuclei on *ESR1* expression. *ESR2* did not significantly differ with treatment or sex, but the POA appeared to have marginally lower expression than the PVN (*p* = 0.05). However, this effect of nuclei did not persist past Benjamin-Hochberg correction (*p_adj_*= 0.21; Supp. Table 4). We did not find any effects of treatment, sex or nuclei on *PRLR* expression (Supp.Table 4).

**Figure 4.**
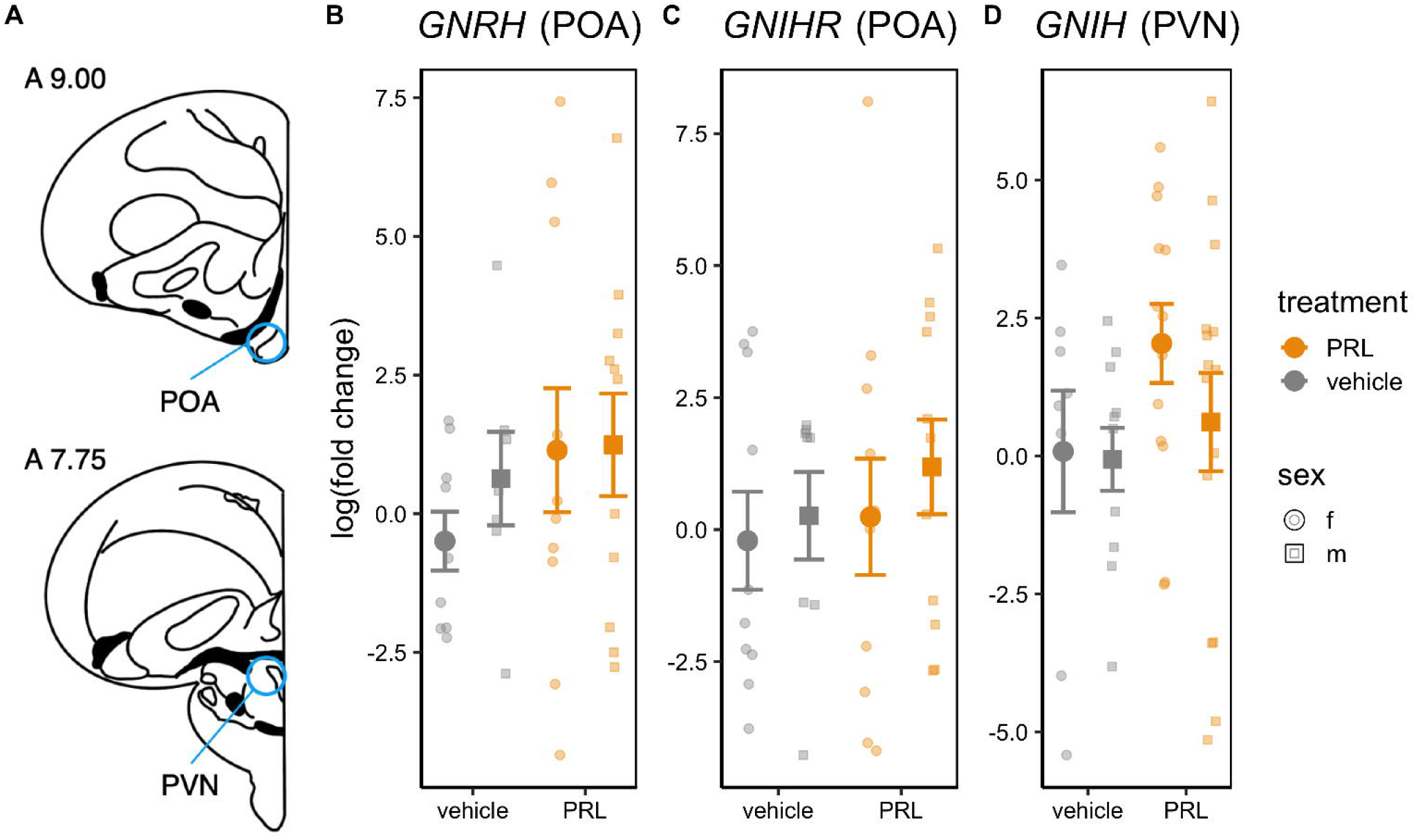
HPG gene expression in hypothalamic nuclei. (A) Genes of interest were measured in microdissected punches of the preoptic area (POA) and paraventricular nucleus (PVN) using the Karstens & Hodos (1966) atlas (representative coronal slices with atlas plate numbers are shown). (B) Gonadotropin releasing hormone (GNRH) expression and ( C) gonadotropin inhibitory-hormone receptor (GnIH-R) expression were measured in the POA, and (D) GnIH itself was measured in the PVN, where GnIH-neuron cell bodies are found. Circles represent females and squares represent males. Mean and SEM are shown for each sex and treatment.

**Table 3.**
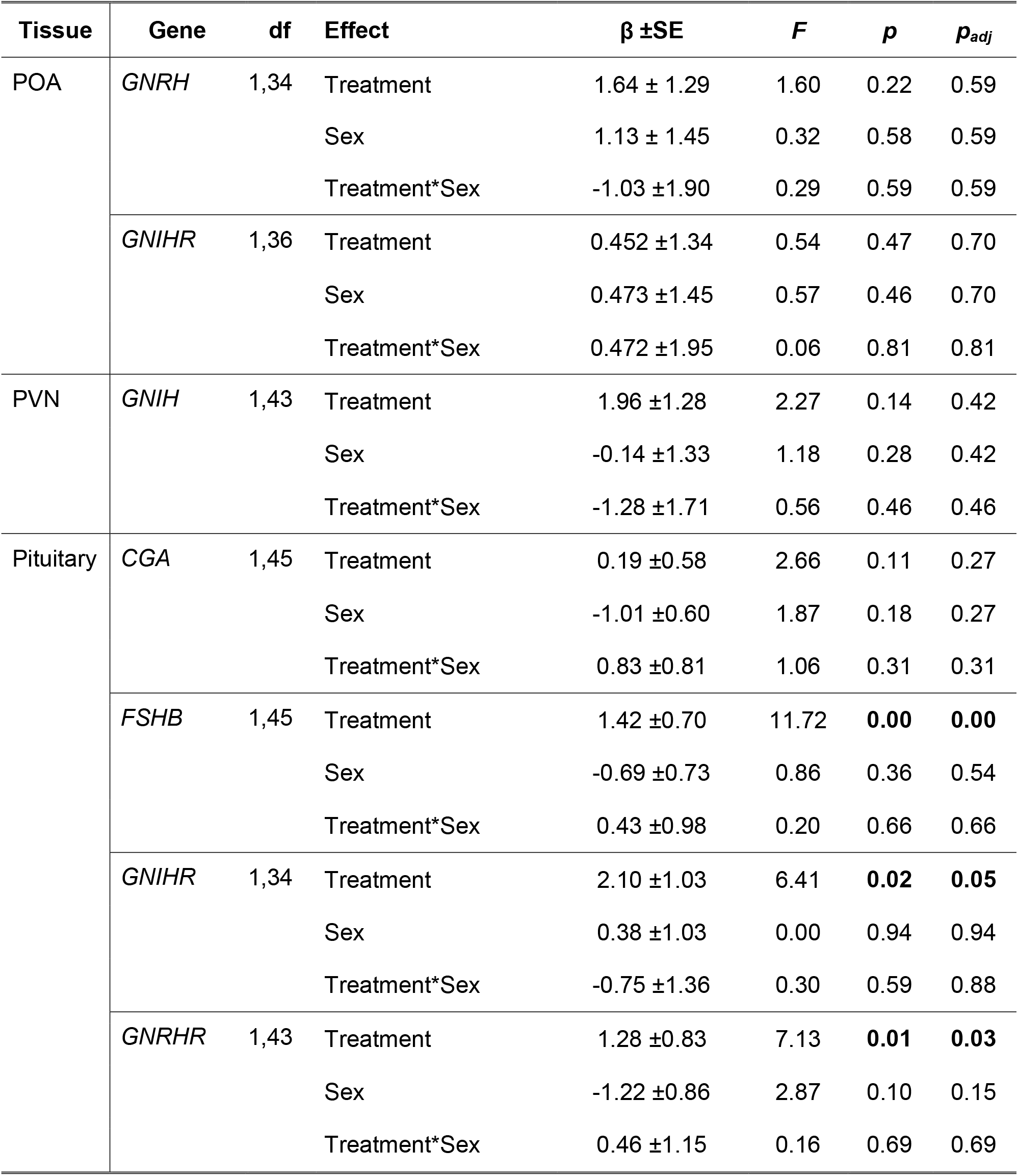
Linear models testing differences in relative gene expression as an effect of treatment, sex, and the interaction for hypothalamic nuclei and the pituitary. Estimates (β) and standard errors are shown in log(fold change), calculated using the Livak & Schmittgen (2001) method. Females treated with vehicle were used as the reference group. Degrees of freedom for each component of the gene linear model. *P*-values are adjusted using the Benjamin-Hochberg false discovery rate correction, significant values (*p* < 0.05) are in bold.

### Pituitary gene expression

In the pituitary, we found a significant effect of treatment on *FSHB* (*p_adj_* < 0.01, Cohen’s *d* = 0.99) but not *CGA* (*p_adj_* > 0.05; Table 3; Fig.5). In terms of receptors, treatment significantly affected *GNRHR* and *GNIHR* expression (Table 3; Fig.5), with oPRL-treated birds expressing higher levels of both receptors than vehicle birds (*GNRHR*, Cohen’s *d* = 0.77 ; *GNIHR*, Cohen’s *d* = 0.85).

**Figure 5.**
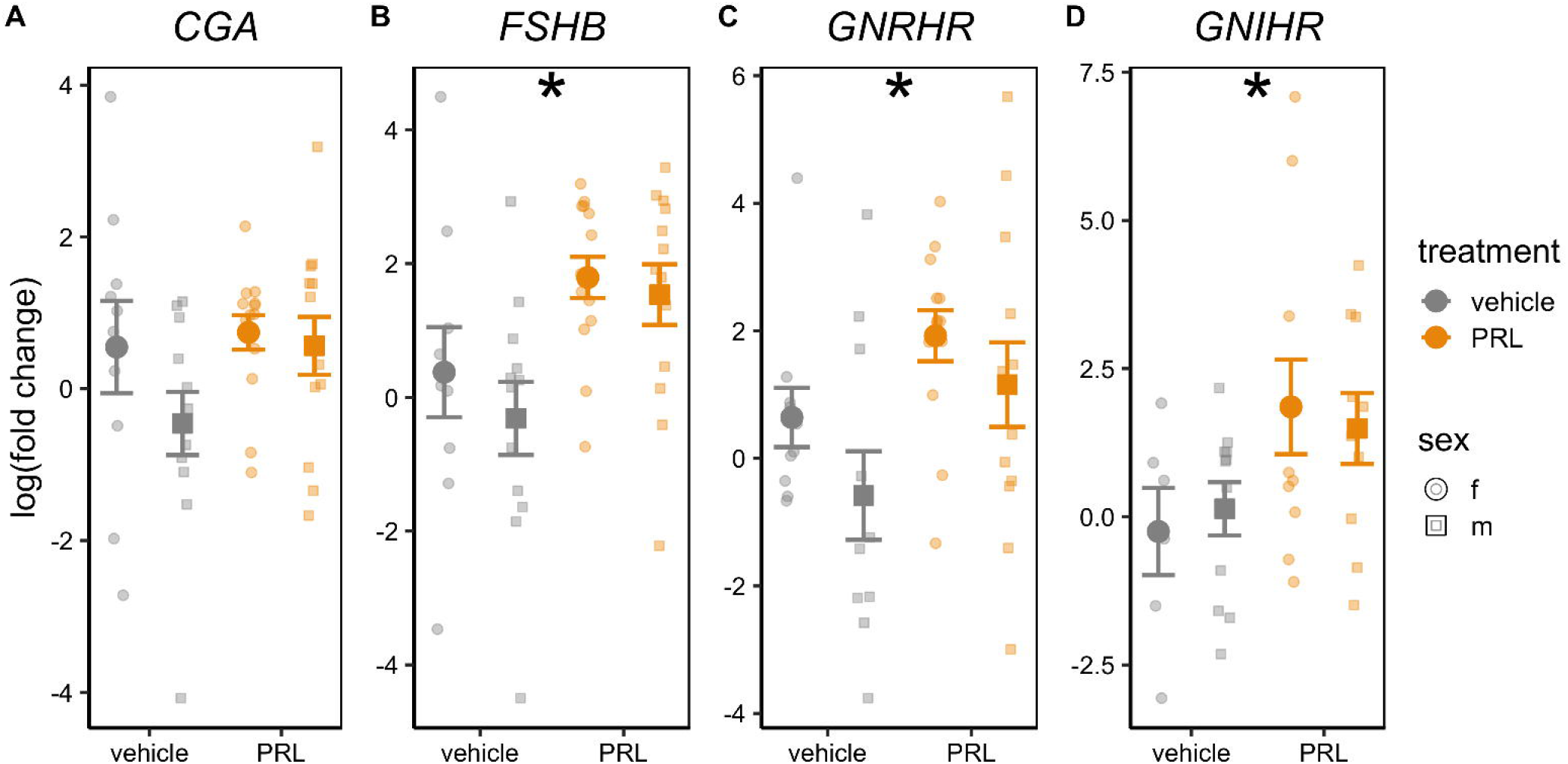
Pituitary gene expression. (**A**) Luteinizing hormone (or CGA), (**B**) follicle stimulating hormone b (FSH), ( **C**) GnRH receptor (GnRH-R) and (**D**) GnIH receptor (GnIH-R) gene expression in the pituitary gland of males and females treated with vehicle or ovine PRL. Mean and SEM are shown for females and males in each treatment group, using circles and squares respectively. Small points represent individual birds. Asterisks (*) represent a significant main effect of treatment (*p_adj_* < 0.05).

### Gonad gene expression

In the testes, we found a significant effect of treatment on *FSHR* and *LHCGR* expression, where oPRL-treated birds expressed higher levels of these receptors than those given vehicle (Table 4; Fig.6A; Cohen’s *d* = 1.49 and 1.41, for *FSHR* and *LHCGR*,respectively). However, treatment had no significant effect on *FSHR* or *LHCGR* in the small white preovulatory follicles in the female ovary(Table 4; Fig.6B). There was no significant effect of treatment on *AROM* or *GNIH* expression in either the testes or the ovaries (Table 4; Fig.6).

**Table 4.**
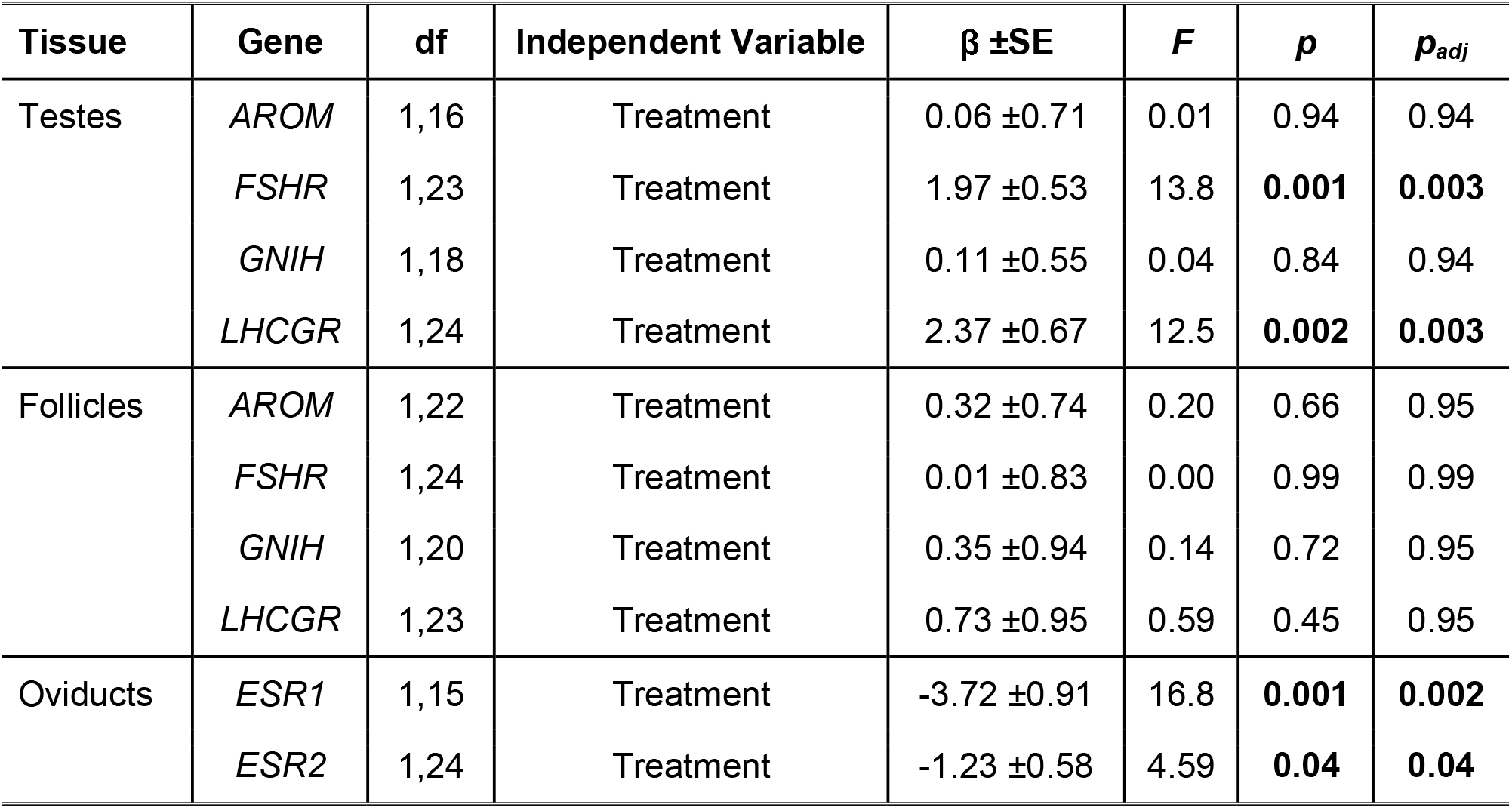
Linear models testing differences in relative gene expression between treatment group in gonadal tissues. Gene expression was measured in testes in males, and small white ovarian follicles and oviducts in females. Estimates (β and standard errors are shown in log(fold change), calculated using the Livak & Schmittgen (2001) method. Vehicle-treated birds are used as a reference group. Degrees of freedom for each component of the gene linear model. P-values are adjusted using the Benjamin- Hochberg false discovery rate correction, significant values (p < 0.05) are in bold.

**Figure 6.**
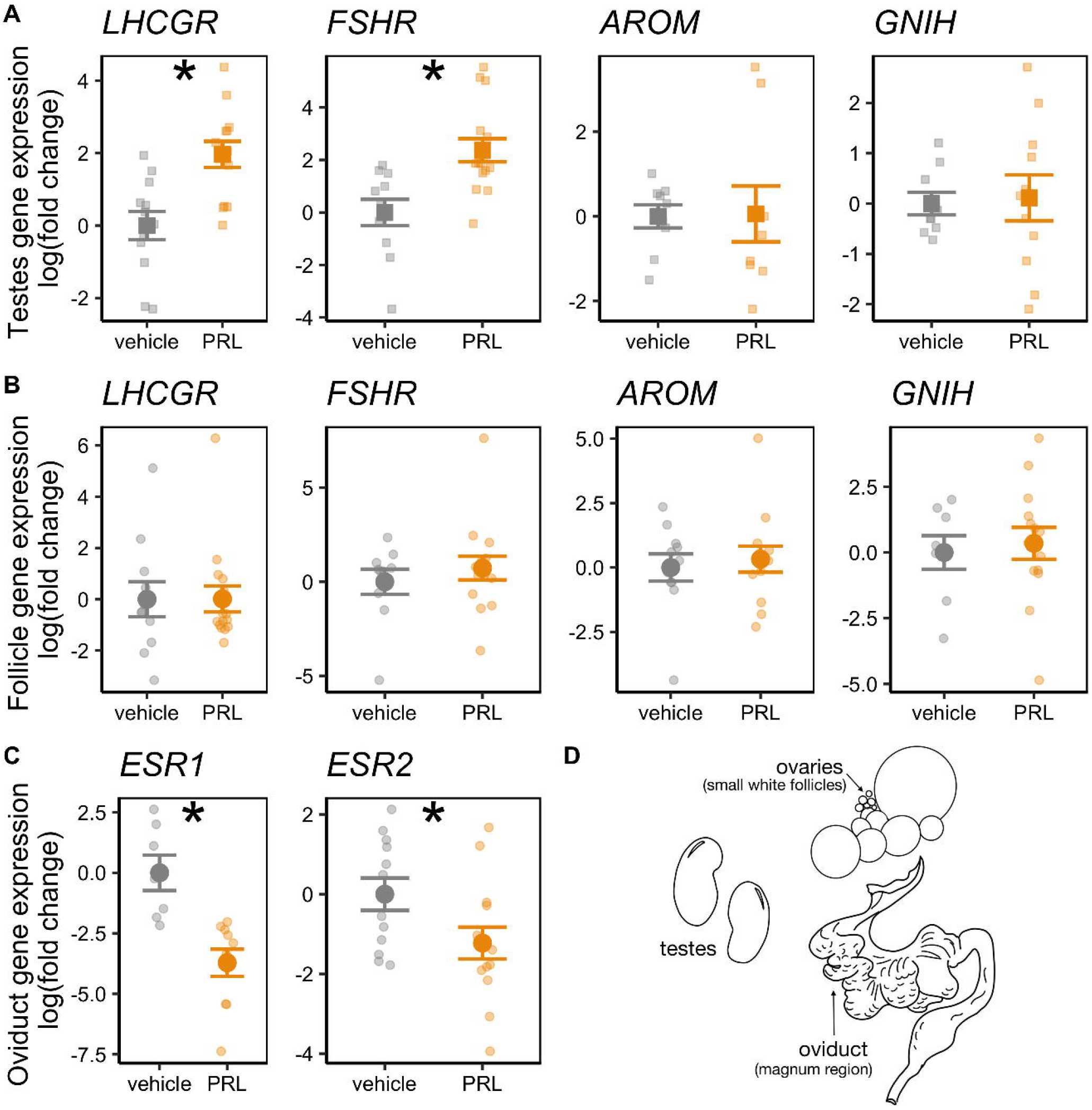
Gonadal gene expression. Relative expression of luteinizing hormone receptor (*LHCGR*), follicle stimulating hormone receptor (*FSHR*), aromatase (*AROM)* and gonadotropin-inhibitory hormone (*GNIH*) relative expression in (**A**) male testes and (**B**)small white ovarian follicles of females , as well as of (**C**) estrogen receptor 1 (*ESR1*) and 2 (*ESR2)* in female oviducts in birds treated with oPRL (orange) or vehicle control (gray). Means and SEM are shown. Small points represent individual birds. Asterisks (*) represent a significant main effect of treatment (*p_adj_* < 0.05). Male tissues are shown with squares and females with circles.

In the oviducts, we found a significant effect of treatment on both estrogen receptors, *ESR1* and *ESR2*, in that oPRL treatment reduced expression of both genes compared to vehicle-treated birds (Table 4; Fig.6C; Cohen’s d = -2.02 and -0.84 for *ESR1* and *ESR2*, respectively).

## Discussion

We found that rock dove pairs given ovine prolactin (oPRL) maintained parental responses after nest loss, but courtship behaviors and copulation rates were not affected compared to vehicle controls.

Hypothalamic gene expression was relatively unaffected by oPRL treatment, but we found increased *FSHB* and hypothalamic hormone receptor expression in the pituitary. The gonads showed sex specific responses, where testes increased in size and gonadotropin receptor expression, while ovaries remained unaffected and oviduct size and estrogen receptor expression decreased. To our knowledge, our study is the first to measure the effect of PRL on courtship and copulation and integrate these behaviors with HPG-wide gene expression.

### Effects on parental and reproductive behaviors

We found that oPRL maintained a parental phenotype, as it increased the likelihood birds would show parental responses towards a novel chick (i.e. feeding, brooding) after nest loss. Supporting this finding, chicks given to oPRL-treated pairs gained weight during the trial (presumably due to regurgitation of crop milk, water or food), whereas chicks in vehicle-treated nests did not. Further, oPRL treatment led to increased crop sac weights, with visible crop milk production, a classic indicator of PRL bioactivity (Lebovic and Nicoll, 1992; Riddle and Braucher, 1931). Together, these results confirm that systemic oPRL administration through the osmotic pumps did bind the avian PRL receptor, leading to crop sac growth and milk production, as well as affected expression of parental behaviors. Our results are consistent with previous studies in ring doves, where systemic PRL administration increased regurgitation feeding rate and chick weight gain in reproductively-experienced doves (Buntin et al., 1991; Wang and Buntin, 1999).

Unlike parental responses, however, courtship and copulation behaviors remained unaffected by PRL treatment. The typical courtship cycle in doves consists of male bow coo to attract their mates’ attention, male nest cooing in potential nest sites, female nest cooing if she accepts the nest site, then both males and females participate in nest building before oviposition (Cheng, 1992; Goodwin, 1983). Pairs cycle through these behaviors in a reliable sequence over a few days (Lehrman, 1964). We found no significant differences between the treatment groups in any of these behaviors, though there was a weak trend suggesting oPRL-treated females may express lower nest cooing. All birds, regardless of treatment, appeared to express courtship behaviors and be at similar courtship stages, with male nest cooing being the dominant behavior for most pairs. We similarly observed no differences in copulation rates. Copulation rates in our study align with those observed in established ring dove pairs allowed to copulate for short periods before ovulation (Cheng et al., 1981). Females in both groups solicited copulations, a behavior coordinated by both GnRH and estrogen actions (Cheng et al., 1981; Gibson and Cheng, 1979). However, female ring doves can exhibit copulatory behaviors even when ovariectomized, showing that ovarian hormones are not necessarily required for this behavior (Cheng, 1973).

The similar courtship and copulation patterns between groups suggests that PRL manipulation did not affect these reproductive behaviors in established pairs. There are several possible explanations. First, our oPRL dose may not have been high enough to inhibit or abolish non-parental reproductive behaviors. Our expected oPRL dose, using calculations described in Sockman et al.(2000), approximates levels seen during mid-chick-rearing in our population (Austin et al., 2021). The mid- to late- chick rearing period can include clutch overlap in rock doves, especially in captive conditions and with experienced pairs (Burley, 1980). Thus, oPRL levels administered may have been similar to a period where mating and courtship co-occur with parental care. Second, stimulatory environmental cues, such as a mate presence, nestbox and nesting material, may have overridden any inhibitory effects of PRL, especially if levels were not supraphysiological. Indeed, mate access and photostimulation lead to nest building and courtship behaviors, as well as increase in gonadotropin release in many bird species, including doves (Cheng and Balthazart, 1982; Lehrman et al., 1961; Shields et al., 1989). Previous studies showing anti-gonadal effects of PRL in doves were conducted in isolation, with birds removed from mates and conspecifics (Buntin et al., 1991, 1988; Foreman et al., 1990). Lastly, courtship and copulation may be relatively unaffected by manipulation of a single hormone in intact animals. Indeed, estradiol implants did not affect courtship and breeding cycle in wild lapland longspurs (Hunt and Wingfield, 2004), and testosterone implants did not affect courtship in captive ring doves (Fusani and Hutchison, 2003). Future studies that manipulate PRL in different doses or in conjunction with other hormones could tease apart these possibilities.

### Effects on hypothalamic gene expression

In the hypothalamus, we found oPRL did not significantly alter GnRH,GnIH, GnRH-R or sex steroid receptor expression. Although not in support of our initial hypothesis, this result is consistent with previous studies in our rock dove population, where hypothalamic gene expression shows fewer significant changes compared to other tissues during parental care (Harris, 2020; Harris et al., 2020). The absence of differences in reproductive behaviors may offer one explanation; mate presence and participation in courtship may have normalized hypothalamic gene expression. For instance, just one to two hours of courtship can increase POA GnRH activity in male doves (Mantei et al., 2008). Our birds interacted continuously for seven days, which may have normalized GnRH expression despite any effects of oPRL. Alternatively, oPRL treatment may have downregulated prolactin responsiveness via *PRLR*, thus reducing the mechanism by which oPRL would alter reproductive gene expression. However, we did not find support for this hypothesis, as *PRLR* did not differ with treatment in either nuclei.

Another possible explanation is that oPRL did not reach the brain or cross the blood brain barrier in our birds given the peripheral placement of osmotic pumps. However, various lines of evidence support that oPRL likely reached the brain. We replicated methods used in the ring dove that showed oPRL treatment led to higher phosphorylated STAT5, a secondary-signalling protein downstream of the PRL receptor, in the PVN and POA (Buntin & Buntin 2014). PRL binding sites have also been well-characterized in the dove brain (Buntin and Ruzycki, 1987; Fechner and Buntin, 1989), and autoradiography experiments show that radiolabeled PRL can cross the blood-brain barrier via the choroid plexus in doves (Buntin and Walsh, 1988) as it does in rodents (Walsh et al., 1987). We also found behavioral differences in parental response that are most likely mediated in part by PRL actions on the brain, though perhaps not through the HPG axis (Grattan and Bridges, 2009).

Despite this evidence, it remains possible that only a portion of the systemic dose reached the brian. Previous studies showing anti-gonadal effects in doves used intracerebroventricular injections to separate out causal neural mechanisms (Buntin et al., 1991, 1988; Buntin and Tesch, 1985). Buntin and Tesch (1985) found that i.c.v. oPRL injections had similar effects on the gonads as the higher doses in previous peripheral administrations (Janik and Buntin, 1985). Even so, not all i.c.v. injections led to gonadal changes or altered gonadotropin release in doves (Foreman et al., 1990). Nonetheless, our methods mimic the physiological release of PRL into the periphery from the pituitary, and we found clear evidence for peripheral effects in our study. At the most conservatively, we can thus interpret PRL actions on the pituitary, gonads and behavior even if only low doses ultimately affected neural gene expression. Direct manipulations of neural PRL are needed to ultimately clarify if the changes in HPG gene expression we observed are mediated through the hypothalamus.

### Effects on pituitary gene expression

In the pituitary, we found significantly increased expression of hypothalamic hormone receptors (*GnRH-R* and *GnIH-R*) and *FSHB* with oPRL treatment. The fact that *FSHB* expression increased, rather than was inhibited, by oPRL was surprising and did not support our original hypothesis. Gonadotropins can be stimulated by salient breeding stimuli, such as presence of nest boxes or a reproductively-receptive mate (Cheng and Balthazart, 1982; Cheng, 1977; Cheng and Follett, 1976; Shields et al., 1989; Silver et al., 1980). In birds, as in mammals, FSH serves to stimulate steroidogenesis and hierarchical development in ovarian follicles (Johnson, 2015) and testes (Deviche et al., 2011). In our study, courtship behaviors were maintained and gonadal size was either maintained or increased with oPRL treatment. Increased *FSHB* expression, combined with increased GnRH receptors, may act as an underlying mechanism to compensate for any potentially inhibitory effects of PRL and maintain reproductive function. Lastly, lack of changes in LH (via *CGA* expression) is consistent with some previous studies, where immunization against PRL or VIP did not affect LH levels (Crisóstomo et al., 1998; Li et al., 2011). LH levels may thus be less sensitive than FSH to PRL manipulation. However, these hypotheses require further testing, as we did not directly measure gonadotropin release.

### Effects on GnIH and GnIH-R expression

A specific aim of this study was to examine the potential relationship between PRL and expression of GnIH and its receptor across the HPG axis. Previous studies show GnIH can increase during transitions in parental care where PRL also rises, such as after hatch or birth in starlings and rodents, respectively (Calisi et al., 2016). While one study shows that GnIH administration does not affect ovine pituitary *PRL* gene expression *in vitro* (Sari et al., 2009), no studies have examined the causal relationship between PRL and GnIH expression to our knowledge. As GnIH inhibits the HPG axis during acute stress and seasonal transitions (Calisi et al., 2011, 2008; Kirby et al., 2009), we hypothesized GnIH may also play a role in any inhibitory effect of PRL on avian reproductive function.

We found that hypothalamic and gonadal *GNIH* expression were unaffected by oPRL treatment, but pituitary *GNIHR* increased in both sexes given oPRL. Classically, hypothalamic GnIH inhibits gonadotropin release by inhibiting GnRH synthesis and release from PVN neurons (Tsutsui and Bentley, 2009; Ubuka et al., 2013), but GnIH can also directly inhibit gonadotropin release by acting on pituitary receptors (Ciccone et al., 2004; Clarke et al., 2008). Locally-expressed gonadal GnIH may also affect reproductive function (Bentley et al., 2017). Our results suggest that oPRL does not affect the HPG axis via hypothalamic or gonadal *GNIH* expression when PRL levels are equivalent to those at mid-chick rearing. Increased *GNIHR* in the pituitary may affect HPG axis regulation (Maddineni et al., 2008), but we did not observe any concordant decrease in gonadotropin expression. It remains possible that PRL manipulation generally dysregulated HPG axis gene expression in a way that would be detrimental to fitness (Bonier and Cox, 2020). However, GnIH neuropeptide translation or release could also have been possibly affected by oPRL treatment, or higher oPRL levels than our dose are required to affect GnIH expression. Further studies examining neural protein levels and/or administering oPRL at doses at the higher end of the physiological range could more definitively determine if the GnIH system is indeed independent of PRL.

### Effects on gonads and gonadal gene expression

In the gonads, we found sex-specific differences in morphology as well as gene expression. We observed slightly *increased* testes size, which stands against the purported “anti-gonadal” role of PRL and contrasts with previous studies in doves (Janik and Buntin, 1985). Design differences may explain this discrepancy; males in our study stayed with their mates in a colonial aviary setting, while birds were isolated in previous studies. Indeed, access to females promotes gonad growth and maintenance in male cowbirds (Dufty and Wingfield, 1986), and spermatogenesis and testosterone release in male starlings (Pinxten et al., 2003; Schwab and Lott, 1969). The presence of mates and the maintenance of courtship behaviors may have led to increased testes size in some males to compensate for any effects of PRL on their own or their mates’ reproductive axis. These differences highlight the influence of environmental stimuli, and the importance of studying hormone regulation in a semi-natural environment (Calisi and Bentley, 2009) in addition to highly-controlled conditions.

Further, gene expression in the testes was consistent with increased testes size. We found oPRL increased *FSHR* and *LHCGR* expression in the testes, suggesting increased responsiveness to gonadotropins in males. LH acts upon Leydig cells to promote androgen synthesis, and FSH stimulates spermatogenesis (Deviche et al., 2011). However, treatments with FSH have been shown to stimulate larger testes growth and development than treatments of LH alone (though both led to growth) (Brown et al., 1975; Deviche et al., 2011). FSH treatment also reduced the inhibitory effect of pharmacological doses of PRL on the testes, and when combined with a lower dose of PRL, actually led to testes growth in non-breeding pigeons (Bates et al., 1937). Taken together with increased pituitary *FSHB* expression, increased *LHCGR* and *FSHR* in oPRL-treated testes is consistent with testes stimulation and the hypothesis that birds may be compensating for any “anti-gonadal effects” of PRL treatment to maintain reproductive behaviors.

In females, oPRL-treatment did not affect ovarian state, but significantly decreased oviduct weight. We found no differences in the largest ovarian follicle diameter between groups, as all birds had a distinct follicular hierarchy with large yolky follicles. PRL has been associated with the end of lay and ovulation in birds (Ryan et al., 2015; Sockman et al., 2000), but other studies show that PRL manipulation does not inhibit or delay laying rates or ovulation (Li et al., 2011; Opel and Proudman, 1984). In ovaries, FSH leads to the development of the follicular hierarchy and release of sex steroids, while LH stimulates ovulation and steroidogenesis (Johnson, 2015; Mishra et al., 2020). The increased pituitary *FSHB* expression we observed may thus have allowed ovarian state to be maintained in oPRL- treated females. Further, ovarian state may have also been maintained by environmental cues in the social aviary setting. As in males, exposure to a mate, courtship and nesting material can also stimulate gonadotropin release and ovarian development in females (Barfield, 1971; Cheng, 1974; Lehrman et al., 1961). Male presence alone can stimulate female doves, as even castrated males, who do not typically show courtship behaviors,can still induce ovulation if prior ovarian development has occurred (Cheng, 1974). Female nest coos themselves can stimulate ovarian activity (Cheng, 1992). As we observed male courtship behaviors and female nest coos in both treatment groups, the interaction of these behaviors and stimuli may have helped maintain ovarian state in the face of oPRL.

Concordantly, we found no differences in gonadotropin receptor or aromatase expression in the small white follicles. PRL can inhibit ovarian steroidogenesis in birds (Camper and Burke, 1977) and in ovaries *in vitro* (Zadworny et al., 1989; but see Hammond et al., 1982; Hrabia et al., 2004). PRL-inhibited steroidogenesis may occur through reduced aromatase expression, specifically in small white follicles (Tabibzadeh et al., 1995); we did not observe this. However, reduced estradiol release and aromatase expression only occurred after 14 days of oPRL treatment in Tabibzadeh et al. 1995, and was not observed after eight days of oPRL (the timeline that aligns more with our study). Additionally, oPRL doses used in these studies may have been higher than ours. Physiological doses of oPRL do not always affect ovarian estradiol and progesterone release *in vitro* (Hammond et al., 1982; Hrabia et al., 2004), suggesting that high doses may be required to inhibit steroidogenesis and affect ovarian gene expression. Although we did not directly measure estradiol levels here, we expect estradiol likely was unaffected given the similar ovarian state and gene expression between treatment groups. However, oPRL may have altered steroidogenesis through other enzymatic pathways and steroid hormone measurements would be required to address this hypothesis.

In contrast, oPRL treatment reduced oviduct size and oviduct estrogen receptor gene expression. Typically, oviduct weight and ovarian follicle state are strongly correlated in birds (Barfield, 1971; Hutchison, 1974). The developed follicular hierarchy releases estradiol, which stimulates the magnum region of the oviduct to grow, differentiate, distend and produce albumin in preparation for laying (Johnson, 2015). Estrogen is more important for this process than gonadotropins, which regulate more of ovarian development (Mishra et al., 2020). Thus, reduced estrogen responsiveness (via reduced *ESR1* and *ESR2* expression) in oPRL-treated females corresponds with observed reductions in oviduct weight. One possibility for the apparent disconnect between ovarian state and oviduct weight may be PRL responsiveness. The oviduct may differ in PRL responsiveness across breeding in a way that the ovaries do not.

Despite recent studies of the oviduct transcriptome (Jeong et al., 2012; Yin et al., 2020), however, links between oviduct gene expression and PRL remain unexplored. Lastly, we cannot rule out the possibility that hormone manipulation disconnected oviduct development from ovarian state, dysregulating the HPG axis in a way that would reduce fitness (Bonier and Cox, 2020).

### Conclusions: Maintaining reproductive state in the face of elevated prolactin

Overall, we found support that PRL manipulation can maintain parental care, but found only mixed support for the hypothesis that PRL acts “anti-gonadally” and inhibits HPG axis regulation. Our results contribute to a mixed literature, where some field and laboratory studies found PRL treatment inhibited gonadal function (Bates et al., 1937; Buntin et al., 1991, 1988, p. 19; Buntin and Tesch, 1985; Janik and Buntin, 1985; Meier, 1969), where others did not (Foreman et al., 1990; Hamner, 1968; Jones, 1969; Li et al., 2011; Meier and Dusseau, 1968; Opel and Proudman, 1984).

One hypothesized reason we did not observe a strong anti-gonadal effect of PRL may be due to the multiple brooding strategy seen in doves and other species. Many birds engage in clutch overlap when resources are available during the breeding season (Burley, 1980; Westmoreland et al., 1986), including seasonal breeders (Grüebler and Naef-Daenzer, 2010; Morton, 2002; Stępniewski and Halupka, 2018; Walsh and Bock, 1997). This reproductive strategy results in co-occurrence of mating (to start a new clutch) and parental care of the still-dependent chicks. A similar phenomenon occurs in some rodents, where females enter a postpartum estrus shortly after birth and lactate pups while pregnant (Connor and Davis, 1980; Roy and Wynne-Edwards, 1995). During multiple brooding, PRL may continue to promote parental behaviors while also occurring at levels low enough to leave gonadal function uninhibited. HPG activity may also naturally increase as PRL declines. In fact, the oPRL dose we used approximates levels observed at nine-days post-hatch in rock doves (Austin et al.,2021), a period where clutch overlap often occurs in experienced pairs in our population (*pers. obs.*). These PRL levels, lower than those seen around late-incubation/hatch, may have allowed both courtship and parental responses to be maintained. Thus, PRL’s effects on the HPG axis appear to depend on the breeding cycle context, even within a species, and this context may be represented by circulating levels. As PRL has often been studied as animals transition out of breeding, such as into seasonal photorefractoriness or molt (e.g., Sharp et al., 1998), subtle shifts in PRL responsiveness *during* breeding may change how this hormone mediates reproductive transitions, especially in species where multiple-brooding occurs.

Taken together, our results support the idea that mating effort and parental care are not mutually-exclusive (Stiver and Alonzo, 2009), and that proximate mechanisms involved in parental care, like prolactin, may not always inhibit other reproductive functions. When we consider contexts where parental and mating behaviors co-occur, such as during multiple brooding, we may find subtle shifts in HPG axis regulation not seen at other breeding stages or in hormone profiles alone. With this in mind, our study highlights the importance of studying hormonal effects on signal production and responsiveness across the entire HPG axis to understand the complex picture of reproductive regulation during breeding transitions.

## Acknowledgements

We thank A.F. Parlow of the National Hormone and Peptide Program at the University of California, Los Angeles for providing us with purified ovine prolactin. K. Hernandez assisted with validating RNA extractions. A. Molina Gil, J. Guerra, A. Kyan, C. Fargeix, A. Quezada, Y. Korplea, J. Arias, and K. Hernandez provided essential animal care and husbandry. T.P.Hahn, B.C. Trainor, R.M.Harris, A.Colon Rodríguez, A. Booth-Griffiths provided helpful discussions and comments.

## Funding

This study was funded by an American Ornithological Society Van Tyne Award (2019) to V.S.F. and NSF IOS grant no.1455960 and CAREER grant no. 1846381 to R.M.C.

## CRediT Authorship Statement

**V.S. Farrar** - Conceptualization, Methodology, Validation, Investigation, Formal Analysis, Writing- Original Draft ; **L. Flores** - Methodology, Validation, Investigation; **R. C. Viernes** - Methodology, Validation, Investigation; **L. Ornelas Pereira** - Investigation; **S. Mushtari** - Investigation; **R.M.Calisi** - Resources, Writing- Review & Editing, Supervision, Funding Acquisition.

## Notes

### Competing Interest Statement

The authors have declared no competing interest.

