## Supplemental Tables 1-4 for "Prolactin maintains parental responses and alters reproductive axis gene expression, but not courtship behaviors, in both sexes of a biparental bird"

#### Supp.Table 1. Hypothalamic nuclei delineations

| **Nuclei** | **Start Plate** | **Start Landmarks** | **End Plate** | **End Landmarks** | **Punch diameter (mm)** |
| --- | --- | --- | --- | --- | --- |
| Preoptic area (POA) | A 9.00 | Tractus septomesencephalicus (TSM) extends to bottom of brain. | A 8.50 | TSM no longer visible, cloudy Tractus quintofrontalis (QF) appears. | 2 |
| Paraventricular nucleus (PVN)* | A 8.25 | QF apparent. | A 6.75 | Tractus opticus (TrO) appears. | 2 |

**Supplementary Table 1. Atlas plates and landmarks used to delineate hypothalamic nuclei.** Plate numbers are referenced from the Karten & Hodos (1966) brain atlas. Landmark names are carried over from the terminology of that atlas. * The PVN is noted as Nucleus paraventricularis magnocellularis (PVM) in Karten & Hodos (1966). For full details on hypothalamic microdissection, see Methods.

#### Supp.Table 2. qPCR Primers

| *Gene (Abbreviation)* | *NCBI Accession number* | *Amplicon length*  *(base pairs)* | | *Efficiency (%)* |  | *Primer sequence* | *Alternative names* |
| --- | --- | --- | --- | --- | --- | --- | --- |
| Androgen receptor (*AR*) | XM_005509361.1 | | 156 |  | F | GCTCTTCTTCAGCATCATTCC |  |
|  |  | |  |  | R | ACCTTGGTGAGCTGGTAAAA |  |
| Aromatase (*AROM*) | XM_021297872.1 | | 125 | 95.6 | F | TGATGATTGCTGCTCCCGAC |  |
|  |  | |  |  | R | ATCTCTGTCACCCATAACAGTCTC | |
| Chorionic gonadotropin alpha (*CGA*) | XM_005499841.2 | | 169 | 99.9 | F | ACAAGGGAGACAGATCATGGA | Equivalent to lutienizing hormone (LH) |
|  |  | |  |  | R | CGCTCCTGGCTTGGAAAAGA |  |
| Estrogen receptor 1 (*ESR1*) | XM_013368702.2 | | 82 |  | F | CCTGTGTCATGTGATCCCTCC | Also known as ER-alpha |
|  |  | |  |  | R | TGGCAGTCCACATTGATCCC |  |
| Estrogen receptor 2 (*ESR2*) | NM_001282841.1 | | 139 | 100.6 | F | GGGAATGATGAAATGTGGCTC | Also known as ER-beta |
|  |  | |  |  | R | GATCTCTTTTACGCGGGTTG |  |
| Follicle stimulating hormone (FSH) | FJ913876.1 | | 140 | 91.2 | F | AGTGAAGATCCCTGGTTGCG |  |
|  |  | |  |  | R | TGAAGGAACAGTAGGACGGC |  |
| Follicle stimulating hormone receptor (*FSHR*) | XM_005498409.3 | | 150 | 97.1 | F | TGCGTGTTTGCTGATGTTCC |  |
|  |  | |  |  | R | AGGACAAATCTCAGTTCGGTGG | |
| Gonadotropin inhibitory hormone (*GNIH*) | XM_005513478.1 | | 144 | 96.3 | F | AAGGTATCACACACAGGCTTGG | Gene name in NCBI: Neuropeptide VF precursor (*NPVF*) |
|  |  | |  |  | R | TAGTCTTCATTTCCCTGGTTCA | |
| Gonadotropin releasing hormone 1 (*GnRH-1*) | XM_005513520.3 | | 105 | 94.1 | F | GAAGTGCAGAAGAGCGAATG |  |
|  |  | |  |  | R | AATCTTCCGTCTGGCTTCTC |  |
| Gonadotropin-releasing hormone receptor (*GNRHR*) | XM_013369955.2 | | 133 | 96.1 | F | GGCACGAGACCCTCTACAAC |  |
|  |  | |  |  | R | TGTGAGGAGAAGAGGCTGGA |  |
| Gonadotropin-inhibitory hormone receptor, or RFamide-related peptide receptor (*GnIHR*) | AB193127.1 | | 119 | 99.4 | F | CTGGACACTGACGCTGCTGA | Also known as RFamide-related peptide receptor (*RFRPR*) or neuropeptide FF receptor 1 (*NPFFR1*) |
|  |  | |  |  | R | GGTTGGCACTGCTGTTGAAG |  |
| Luteinzing hormone choriogonadotropic hormone receptor (*LHCGR*) | XM_021287698.1 | | 167 | 90 | F | CCAGATGTCCTGGATGTTTCTT | Also known as Lutropin-choriogonadotropic hormone receptor |
|  |  | |  |  | R | CAGTGGCTGGGATACGTTAGA | |
| Prolactin receptor *(PRLR*) | NM_001282822.1 | | 158 | 95.2 | F | TCTTCCTTGCACACATGAAACC | |
|  |  | |  |  | R | TCCAGGGTATGATTGACCAGT | |
| *Reference genes* | | | | | | | |
| Beta actin (*ACTB*) | XM_005504502.2 | | 107 | 95.8 | F | ATGTGGATCAGCAAGCAGGAG | |
|  |  | |  |  | R | CATTTCATCACAAGGGTGTGGG | |
| Glyceraldehyde 3-phosphate dehydrogenase (*GAPDH*) | NM_001282835.1 | | 99 |  | F | AGCAATGCTTCCTGCACTAC |  |
|  |  | |  |  | R | CTGTCTTCTGTGTGGCTGTG |  |
| Hypoxanthine phosphoribosyltransferase 1 (*HPRT1*) | XM_005500563.2 | | 150 | 94.72 | F | GCCCCATCGTCATACGCTTT |  |
|  |  | |  |  | R | GGGGCAGCAATAGTCGGTAG |  |
| Ribosomal protein L4 (*RPL4*) | XM_005511196.1 | | 78 | 105.4 | F | GCCGGAAAGGGCAAAATGAG |  |
|  |  | |  |  | R | GCCGTTGTCCTCGTTGTAGA |  |

**Supplemental Table 2. Primers used in quantitative PCR.** Gene names and abbreviations, and NCBI GenBank accession numbers used to design primers are listed. The amplicon length is shown in base pairs, primer efficiency, and forward and reverse primers sequence is shown for each validated primer. Alternative names, including those used in GenBank or other citations, are also listed.

#### Supp.Table 3. Reference gene stability

|  | POA | PVN | Pituitary | Testes | Ovarian Follicles | Oviducts |
| --- | --- | --- | --- | --- | --- | --- |
| Treatment | 1.14 (0.292) | 1.57  (0.216) | 0.04  (0.851) | 1.01  (0.323) | 1.21  (0.282) | 0.10  (0.944) |
| Sex | 0.24  (0.627) | 1.76  (0.191) | 0.40  (0.530) |  |  |  |
| Treatment*Sex | 0.13  (0.718) | 0.184  (0.670) | 0.58  (0.449) |  |  |  |

#### **Supplemental Table 3**. **Reference gene stability across treatment, sex, and their interaction for each tissue**. For each tissue, we ran ANOVA upon general linear models of the form: *mean reference gene Ct ~ treatment*sex*, except for gonadal tissues and oviducts, where only treatment was included as an independent variable. *F* statistics from these ANOVA for each factor are shown, along with *p* values in parentheses. No reference gene combination showed significant differences across treatment, sex, or their interaction for any tissue, illustrating that reference genes were indeed stable in these tissues.

#### Supp.Table 4. Hyp Sex Steroid lms

Sex Steroid and Prolactin Receptors (Measured in both nuclei)

| ***Tissue*** | ***Gene*** | ***Independent Variable*** | ***β ± SE*** | ***F*** | ***p*** | ***p_adj_*** |
| --- | --- | --- | --- | --- | --- | --- |
| Hypothalamus  (POA & PVN) | *AR* | Treatment | -0.89 ± 0.81 | 1.41 | 0.24 | 0.61 |
|  |  | Sex | -0.55 ± 0.81 | 0.27 | 0.61 | 0.61 |
|  |  | Nuclei | -0.50 ± 0.62 | 0.70 | 0.41 | 0.61 |
|  |  | Treatment*Sex | 0.56 ± 1.07 | 0.27 | 0.61 | 0.61 |
|  | *ESR1* | Treatment | -0.79 ± 0.83 | 4.49 | 0.04 | 0.16 |
|  |  | Sex | 0.37 ± 0.84 | 0.00 | 0.99 | 0.99 |
|  |  | Nuclei | -0.52 ± 0.66 | 0.60 | 0.44 | 0.76 |
|  |  | Treatment*Sex | -0.65 ± 1.13 | 0.33 | 0.57 | 0.76 |
|  | *ESR2* | Treatment | -0.39 ± 0.65 | 0.76 | 0.38 | 0.77 |
|  |  | Sex | -0.001 ± 0.68 | 0.001 | 0.98 | 0.98 |
|  |  | Nuclei | -0.86 ± 0.44 | 3.87 | 0.05 | 0.21 |
|  |  | Treatment*Sex | 0.05 ± 0.89 | 0.003 | 0.95 | 0.98 |
|  | *PRLR* | Treatment | -1.21 ± 0.51 | 1.46 | 0.23 | 0.76 |
|  |  | Sex | 0.88 ± 0.52 | 0.79 | 0.38 | 0.76 |
|  |  | Nuclei | 0.08 ± 0.40 | 0.01 | 0.93 | 0.93 |
|  |  | Treatment*Sex | -0.08 ± 0.69 | 0.01 | 0.93 | 0.93 |
